# Flood-induced interspecific interactions in spring-fed tributary as an ecosystem function of heterogeneous river networks

**DOI:** 10.1101/2020.04.26.062968

**Authors:** Masaru Sakai, Ryoshiro Wakiya, Gosuke Hoshi

## Abstract

Understanding the migratory community dynamics of river networks is important for maintaining lotic system integrity. River animals migrate to their preferred habitats in spatiotemporally heterogeneous river environments. Spring-fed habitats are uniquely characterized by stable temperature and flow regimes, which create suitable spawning habitats for the chum salmon *Oncorhynchus keta. O. keta* exhibits “run up” to its birthplace for spawning, especially during floods. Because the eggs deposited by this anadromous fish are nutritious and actively consumed by freshwater animals, the location and timing of *O. keta* spawning events affect the spatiotemporal accumulation of mobile consumers. In this study, we examined changes in temporal population density in spawning *O. keta* and a mobile consumer (juvenile *O. masou masou*) in a lowland, spring-fed tributary in northern Japan during a 48.5-mm autumn rainfall event. In both species, population density increased, and then decreased, after the rainfall event. In *O. keta*, these changes were closely associated with rainfall intensity, whereas in *O. masou masou* the peak was delayed until 3 days after the rainfall event. A comparison of the gut contents of *O. masou masou* sampled from a spring-fed tributary and an adjacent non-spring-fed tributary indicated greater consumption of *O. keta* eggs in the spring-fed tributary. These results suggested that preferential migration of *O. keta* into spring-fed tributaries for spawning induces subsequent accumulation of juvenile *O. masou masou*, in turn increasing *O. keta* egg consumption. These findings improve our understanding of community dynamics during floods in a heterogeneous river network environment.

## Introduction

Spring-fed streams, which are maintained by continuous groundwater upwelling, constitute unique ecosystems with stable temperature and flow regimes (Mattson et al. 1995; Sear et al. 1999; Lusardi et al. 2016). The narrow range of water temperature in spring habitats creates refugia for stenothermal glacial relict species (Reiss et al. 2016; Sun et al. 2020). Stable flow regimes in spring-fed streams are associated with less sediment runoff during floods than non-spring-fed streams, and the formation of fine substrates inhabited by abundant detritivorous macroinvertebrates (Sakai et al. 2020). Thus, the unique habitats of spring-fed streams enhance beta diversity in river networks (Reiss et al. 2016; Sakai et al. 2020).

The chum salmon *Oncorhynchus keta*, an important North Pacific fishery species, is an anadromous fish. After 2–4 years of growth in the ocean, *O. keta* exhibits “run up” to its birthplace for spawning from summer to winter (Masuda et al. 1984), especially during floods caused by rainfall events (Banks 1969). *O. keta* has preferentially selected upwelling water habitats with a stable warm temperature for spawning (Kobayashi 1968; Milligan et al. 1984; Geist et al. 2011); these upwelling springs form egg accumulation zones after rainfall events during the spawning season.

*O. keta* eggs are a significant and nutritious food resource for freshwater fishes. For example, juvenile salmonids rely heavily on eggs deposited by anadromous spawning salmon, including *O. keta* (Armstrong et al. 2013; Koshino et al. 2013; Bailey et al. 2019). This marine-derived resource subsidy is available both after floods in spring habitats, and spatiotemporal heterogeneity in its availability influences the migration of mobile consumers in rivers containing spawning salmon species (e.g., Armstrong et al. 2013).

Understanding migratory community dynamics in river networks is important to preserve lotic system integrity (Ward et al. 1999; Elosegi et al. 2010); however, direct observation of the migration of lotic animals, particularly fishes, during floods is generally difficult and unsafe. In contrast, spring-fed streams are suitable for investigating community dynamics during flood events due to their high stability. Autumn rainfall can trigger upstream migration of *O. keta* for spawning, which influences the foraging habitats of species that feed on their eggs. This interactional process may represent an additional ecosystem function of spring-fed streams, i.e., structuring biota in river networks. However, substantial gaps in understanding remain regarding the functional role of spring-fed streams in the dynamics of stream fishes during floods.

Because springs generally provide purified water, spring-fed streams are frequently reserved for use as a human water supply (LaMoreaux and Tanner 2001). Elucidating the ecosystem functions of spring-fed streams would further highlight their value. In the present study, we investigated temporal variation in stream fish dynamics in a spring-fed tributary after a rainfall event. Although access to the surrounding non-spring-fed streams was not feasible during the rainfall event, the spring-fed tributary was relatively calm and could be safely accessed for surveys. We examined the responses of spawning *O. keta* and juvenile *O. masou masou* to the flood event, and associated interspecific interactions related to the consumption of *O. keta* eggs. The findings of this study contribute to our understanding of community dynamics during floods in heterogeneous river network environments.

## Materials and methods

### Study site

This study was conducted in the Shubuto River basin in Kuromatsunai, Hokkaido, Japan (42.64° N, 140.34° E), which encompasses an area of 367 km^2^ and includes both montane and lowland regions. The region is underlain by sandstone and mudstone, including Cenozoic fossil shells and tuff. The mean annual precipitation between 2009 and 2018 was 1,615.8 mm, and the mean air temperature was 7.5°C, measured at the Kuromatsunai AMeDAS automated weather station located 4 km northwest of the study site. The dominant tree species in riparian zones within the study area are *Salix* species and *Quercus crispula*, and the dominant understory plants are *Sasa kurilensis* and *Reynoutria sachalinensis*. No dams or weirs prevent the migration of fishes in the mainstem of the Shubuto River (Miyazaki and Terui 2016). The Shubuto River system has a perennial spring-fed tributary with significant groundwater discharge (ca. 0.1 m^3^/s) in the lowland region (Kamiyama River, Fig. 1). This stream has no artificial structures that inhibit longitudinal connections, and stream fishes thus migrate between the spring-fed tributary and mainstem.

**Fig. 1.**
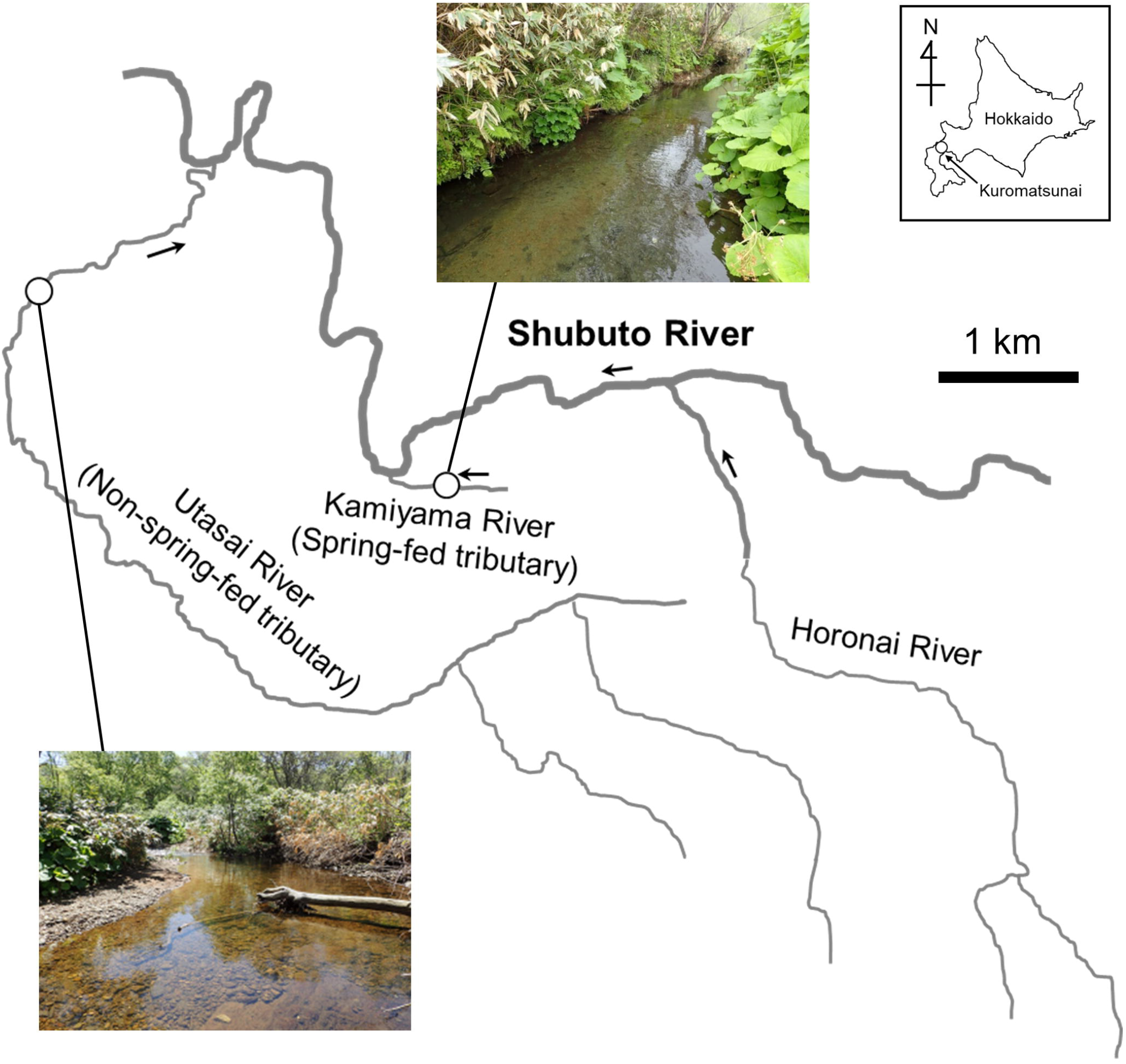
Map and photographs of the study tributaries.

At this spring-fed tributary, we examined temporal changes in the population density of two *Oncorhynchus* fishes (*O. keta* and *O. masou masou*) during a rainfall event. On October 6–7, 2018, a temperate low-pressure system (formerly the Kong-rey typhoon) released a total of 48.5 mm of rain at the study site. According to our visual surveys performed immediately after the rainfall event, water levels increased substantially in most non-spring-fed streams of the Shubuto River system; however, the study tributary showed no dramatic increase in water level. We collected fish both before (October 6) and after (October 7, 9, and 10) the rainfall event; hereafter, we refer to these collection dates as “before,” “after d1,” “after d3,” and “after d4”, respectively.

We also collected the gut contents of *O. masou masou* in an adjacent perennial non-spring-fed tributary (Utasai River, Fig. 1), for comparison between spring-fed and non-spring-fed tributaries after the rainfall event. Stream width and water depth were similar between the tributaries (Table 1); however, the non-spring-fed tributary had a gravel-dominated streambed with a faster current. *O. keta* rarely migrates to the non-spring-fed tributary during spawning seasons, but migrates yearly to the spring-fed tributary; a preference for spring-fed habitats as spawning substrates in this species has been reported previously (e.g., Kobayashi 1968; Milligan et al. 1986), leading to the construction of a facility for capturing mature *O. keta* in this tributary. The facility irreversibly traps mature *O. keta* in an artificial pool, and trapped individuals are transported to another tributary of the Shubuto River system for egg collection and fry rearing. Thus, eggs found in the spring-fed tributary are presumed to have spawned within the tributary, and not drifted from the facility. On October 10 (after d4), when both tributaries had ordinary flow conditions, we sampled *O. masou masou* gut contents.

**Table 1.** Mean ± 1 standard deviation of environmental variables in the spring-fed and non-spring-fed tributaries.

| Variable | Spring-fed tributary |  |  | Non-spring-fed tributary |  |  |
| --- | --- | --- | --- | --- | --- | --- |
| | Mean | ± | SD | Mean | ± | SD |
| Stream width (m) | 4.07 | ± | 0.25 | 5.13 | ± | 0.91 |
| Water depth (cm) | 15.02 | ± | 3.60 | 16.64 | ± | 4.99 |
| Current velocity (cm/s) | 12.43 | ± | 7.09 | 23.72 | ± | 10.35 |
| Fine sediment deposition (%)* | 80.63 | ± | 9.81 | 22.50 | ± | 15.92 |

\*cover of fine sediment (< 2 mm) on streambed estimated using the method in Sakai et al. (2013).

### Hydrological monitoring

Water level and temperature loggers (HOBO CO-U20L-04; Onset, Bourne, MA, USA) were fixed on streambeds of the tributaries using metal tubes and wire, and hydraulic pressure was recorded once per hour. Before the rainfall event started, we preliminarily recorded water levels at the monitoring points on October 6–7, 2018 using a folding scale, and devised linear regression models of the relationship between water level (cm) and hydraulic pressure (kPa). Based on these models, hydraulic pressure data recorded in the loggers were transformed into water level data. For both study reaches, the water level at 1:00 A.M. on October 6, 2018 was set at 0 cm, and hydrographs of the ensuing water level changes were created. Hourly rainfall data were obtained from the Kuromatsunai AMeDAS automated weather station.

### Fish collection

A 40-m-long study reach was established in the spring-fed tributary to investigate temporal changes in *O. keta* and *O. masou masou* population density during a rainfall event. The Shubuto River *O. masou masou* population includes both anadromous and resident individuals; most females and some males become anadromous after approximately 2 years of growth in the river (Miyazaki 2017). Thus, small individuals collected in the study area were mainly juveniles and resident males. Because the rainfall event occurred during the reproductive period, we classified small *O*. *masou masou* individuals (standard body length = 60–160 mm) as mature or juvenile based on the presence or absence of nuptial coloration. *O. keta* is anadromous, and generally runs up to the Shubuto River from mid-September, where it spawns until mid-November after 2–8 years of growth in the North Pacific (Miyazaki 2017). Therefore, individuals found in the river during the autumn reproduction season are mature.

We conducted electrofishing using a backpack electrofisher (LR-20; Smith-Root Inc., Vancouver, WA, USA). The upper and lower limits of the tributary study reach were gently partitioned using fishnets to prevent fishes from entering or exiting the reach during collection. Three-pass electrofishing was conducted from lower to upper positions, to collect as many *Oncorhynchus* fish as possible. After collection, the wetted width was measured at five points (10-m intervals) within the study reach, to better represent the surveyed area during each collection (before, after d1, d3, and d4) allow for accurate calculation of fish population density.

The population density of *O. masou masou* was estimated using the maximum likelihood method proposed by Zippin (1956), because the number of collected individuals decreased substantially between the first and third removals. The number of collected *O. keta* individuals varied and exhibited no clear pattern. However, we confirmed that each three-pass removal collected all individuals within the reach; this was possible due to their large body size (standard length = 425–801 mm). Therefore, we used the total number of collected individuals to directly estimate population density for this species. The maximum likelihood method was applied using R software (v. 3.6.3; R Core Team 2020) with the *FSA* package (Ogle et al. 2020).

### Gut content collection

In the early morning and evening of after d4, we collected *O. masou masou* for gut content analysis using a stomach pump; a total of 31 and 11 individuals were collected in the spring-fed and non-spring-fed tributaries, respectively. Each gut content sample was immediately preserved in 70% ethanol. The body size distribution of the collected *O. masou masou* was similar between tributaries. Fish were released into their original habitats after gut content collection. To avoid interference with our population density estimates in the spring-fed tributary, we performed population density sampling in this tributary in the early morning of after d4, prior to gut content collection.

Gut contents were sorted and identified using a stereomicroscope (SZ61; Olympus, Tokyo, Japan). Identified gut contents were dried in an oven (FC-610; Advantec, Tokyo, Japan) at 60°C for 24 h and then weighted precisely to within 0.01 mg. To test for differences in the gut contents of *O. masou masou* between tributaries, we analyzed the dry weight data using one-way analysis of variance (ANOVA) with permutation tests, which allowed us to deal with unbalanced sample sizes. ANOVA was performed using the R 3.6.3 software with the *anova.1way* function (Legendre 2007).

## Results

Following the onset of the rainfall event, the water level in the spring-fed tributary declined slightly, but was clearly lower overall than that in the non-spring-fed tributary throughout the monitoring period (Fig. 2). Water levels in the non-spring-fed tributary reflected changes in rainfall intensity, and peaked at 29 cm above the initial water level (Fig. 2). A total of 175 individuals of *O. masou masou* and 35 individuals of *O. keta* were collected in the spring-fed tributary. In both species, population density increased after the rainfall event, but subsequently decreased (Fig. 3); that of *O*. *masou masou* peaked at after d3, whereas for *O. keta* it decreased after d1. All collected *O*. *masou masou* were juveniles, and all *O. keta* individuals were mature and reproductive.

**Fig. 2.**
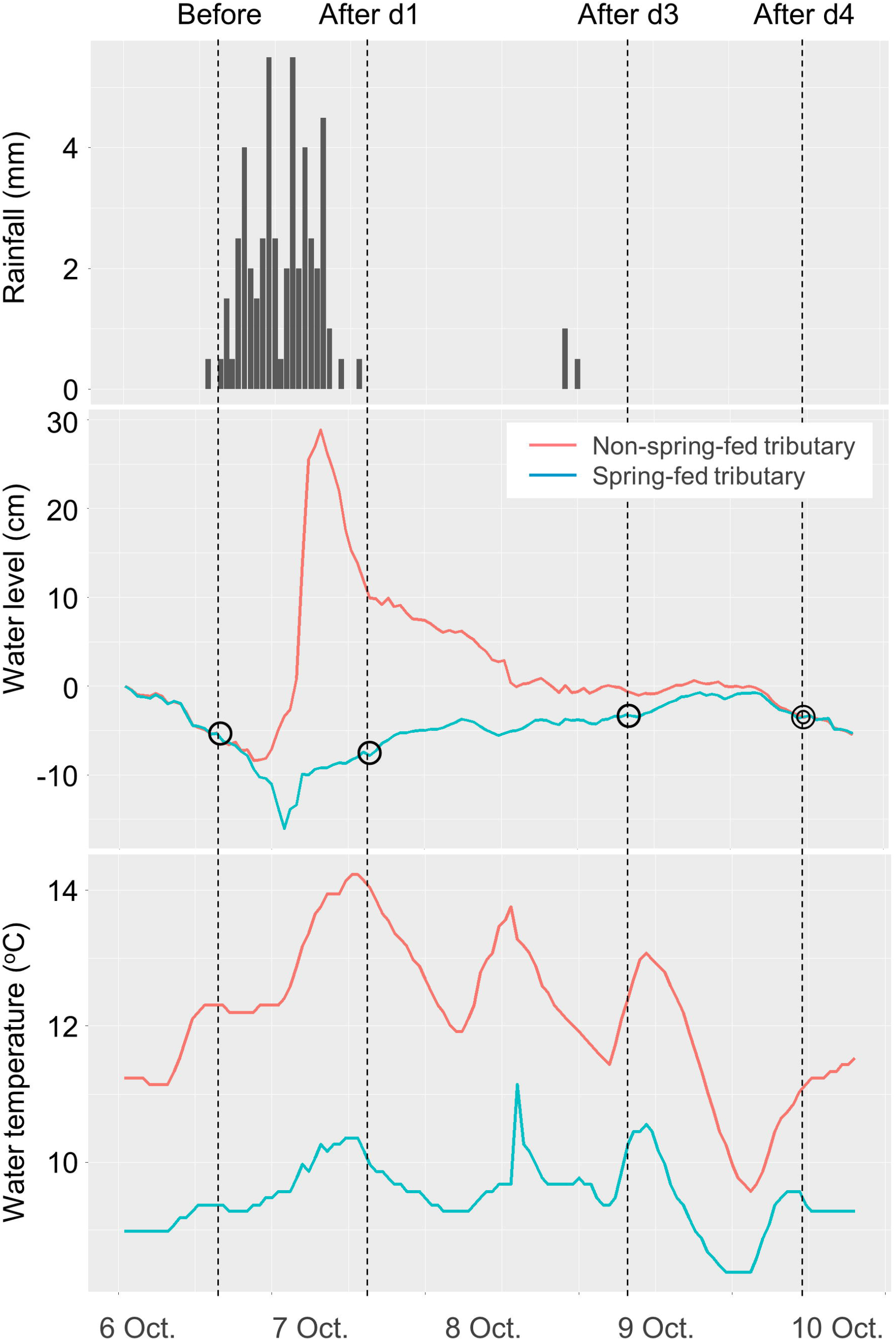
Hydrographs and water temperature data for spring-fed and non-spring-fed tributaries during October 6–10, 2018. Circles indicate dates of fish collection for population density estimates in the spring-fed tributary; double circle indicates the date of additional fish collection in both tributaries for gut content analysis.

**Fig. 3.**
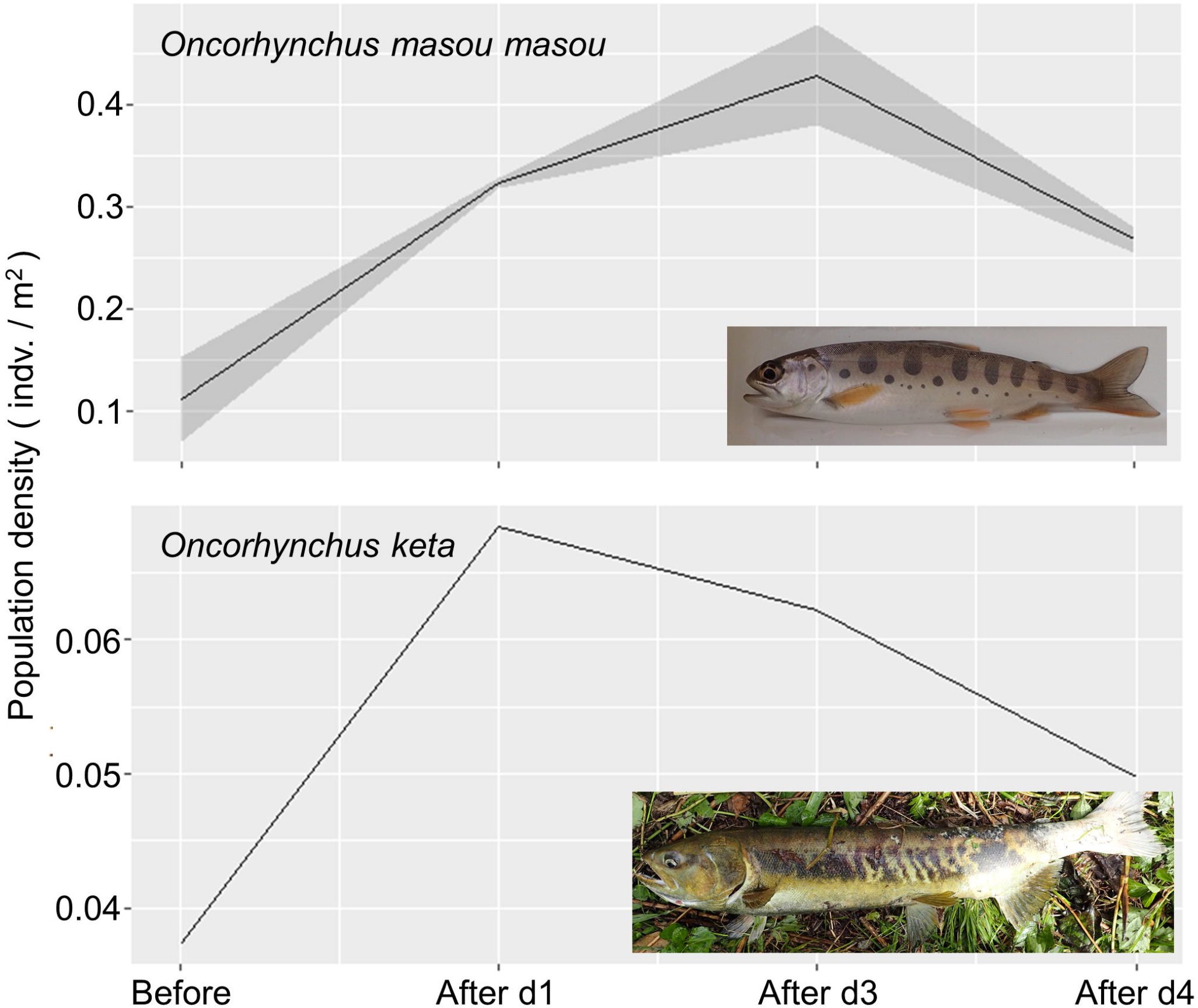
Temporal changes in the population density of *Oncorhynchus masou masou* and *O. keta* in the spring-fed tributary. Gray band indicates the 95% confidence interval for *O. masou masou*.

At after d4, *O. masou masou* in the spring-fed tributary consumed significantly more *O. keta* eggs, whereas those in the non-spring-fed tributary consumed more hemipteran, hymenopteran, and trichopteran insects (Table 2). Due to the large contribution of *O. keta* eggs, the total gut contents were higher in the spring-fed tributary than in the non-spring-fed tributary (Table 2).

**Table 2.** Mean ± 1 standard deviation of dry weight (mg) of gut contents in each order and total in the spring-fed and non-spring-fed tributaries.

|  | Spring-fed tributary |  |  | Non-spring-fed tributary |  |  | *F* | *P* |
| --- | --- | --- | --- | --- | --- | --- | --- | --- |
| | Mean | ± | SD | Mean | ± | SD | | |
| Amphipoda | 0.52 | ± | 1.04 | 0.00 | ± | 0.00 | 2.63 | 0.14 |
| Acari | 0.01 | ± | 0.03 | 0.00 | ± | 0.00 | 0.98 | 0.35 |
| Ephemeroptera | 0.86 | ± | 2.72 | 0.68 | ± | 0.68 | 0.05 | 0.82 |
| Plecoptera | 0.01 | ± | 0.04 | 0.02 | ± | 0.06 | 0.49 | 1.00 |
| Hemiptera | 0.00 | ± | 0.00 | 1.78 | ± | 3.75 | 7.34 | **0.01** |
| Hymenoptera | 0.00 | ± | 0.00 | 0.56 | ± | 1.04 | 9.27 | **0.02** |
| Coleoptera | 0.00 | ± | 0.00 | 0.03 | ± | 0.11 | 0.09 | 0.25 |
| Trichoptera | 0.27 | ± | 1.07 | 1.35 | ± | 1.03 | 8.44 | **0.01** |
| Diptera | 1.22 | ± | 2.04 | 0.75 | ± | 0.70 | 0.54 | 0.45 |
| Salmoniformes | 9.30 | ± | 12.78 | 0.00 | ± | 0.00 | 5.73 | **0.02** |
| Total | 12.18 | ± | 13.72 | 5.17 | ± | 4.56 | 2.72 | 0.10 |

Bold characters indicate statistically significant difference between the tributaries, according to one-way analyses of variances with permutation tests ( $P < 0.05$ ).

## Discussion

The results showed that increase in population density of *O. keta* immediately after the rainfall event, and the subsequent peak of population density of juvenile *O. masou masou* was observed three days after the rainfall event in the spring-fed tributary. The individuals of *O. masou masou* in the spring-fed tributary consumed more eggs of *O. keta* compared to those in the adjacent non-spring-fed tributary. These results suggest that preferential migration of *O. keta* into spring-fed tributaries for spawning induces subsequent accumulation of juvenile *O. masou masou*, in turn increasing *O. keta* egg consumption.

Typically, lowland spring-fed streams formed by groundwater discharge are not deep (Sear et al. 1999), and their flow regimes are generally stable (Lusardi et al. 2016). In this study, water levels in the spring-fed tributary did not exhibit the same marked increase observed in the non-spring-fed tributary during the rainfall event. This stability creates a valuable opportunity for direct field observation of the dynamics of aquatic animal communities during floods.

Hydrological monitoring indicated that the water level of the spring-fed tributary decreased following the onset of rainfall. This change may be attributable to the onset of water transmission into the *O*. *keta* egg collection facility, because local fishermen know that this species actively migrates to upstream spawning habitats during floods (Banks 1969). Our results also indicated that the population density of *O. keta* peaked immediately after the rainfall intensity peak. The dependence of *O. keta* on spring-fed habitats for spawning (Kobayashi 1968; Milligan et al. 1984; Geist et al. 2011) may explain the accumulation of this species in the spring-fed tributary during the flood.

Although the population density of *O. keta* was sensitive to temporal changes in rainfall intensity, the peak population density of *O. masou masou* was delayed until 3 days after the flood. We presume that this temporal increase in *O. masou masou* population density was influenced by two factors: evacuation to flow refugia and acquisition of *O. keta* eggs as food. Tributaries have been considered as flow refugia for fishes (Koizumi et al. 2013); thus, some juvenile *O. masou masou* may migrate to spring-fed tributaries to avoid floods. However, the study reach is located 500 m upstream of the confluence of the tributary and the mainstem; this distance may be too long for such evacuations. Makiguchi et al. (2009) reported that a related species, *O. masou formosanus*, endured a massive typhoon event, which induced a 3-m water level increase, by remaining within their original habitats in the mainstem. Therefore, the moderate flood observed in this study may not have been a major driver of fish evacuation to the spring-fed tributary.

The eggs of anadromous salmon species are a significant and nutritious food resource for freshwater fishes (Armstrong et al. 2013; Koshino et al. 2013; Bailey et al. 2019). This food resource is likely targeted by juvenile *O. masou masou* accumulating in the spring-fed tributary. Our gut content analysis indicated that there were significantly more salmon eggs in the spring-fed tributary than in the adjacent non-spring-fed tributary, where *O. keta* rarely migrates. The time lag between the population density peaks of *O. keta* and *O. masou masou* may be attributable to differences in their swimming abilities and/or the detection of egg production cues by *O. masou masou*, perhaps related to the timing of *O. keta* sperm release and downward dispersion. Because the present study lacks information on the redd distribution of *O. keta* in the study streams before and after the rainfall event, whether the consumed eggs by *O. masou masou* were spawned after the rainfall event or not was unknown. Grasping the redd distribution can help explain the plausible process proposed here more clearly.

Both spring-fed streams and upwelling water habitats function as spawning habitats for *O. keta* (Kobayashi 1968; Milligan et al. 1984; Geist et al. 2011); accumulation of *O. keta* in such habitats can occur during floods (Banks 1969). In this study, we directly observed upward migration of *O. keta* into the spring-fed tributary due to its stable flow regime. Our results indicate that the ecosystem functions of spring-fed streams, with respect to the formation of a unique habitat with stable temperature, provide mature *O. keta* with a suitable habitat and, in turn, provide other species with marine-derived food resources. Interspecific interactions with juvenile *O. masou masou* also occur in such habitats in spring. Thus, it is crucial to evaluate community dynamics in river networks in spatiotemporally heterogeneous environments (Armstrong et al. 2013). A robust sampling design with replications for tributaries and rainfall events, lacked in this study, can unravel the ecosystem function of heterogeneous river networks more clearly. The findings of the present study highlight the importance of enhancing spatiotemporal heterogeneity in spring-fed habitats to maintain interspecific interactions, which have implications for freshwater biodiversity and resource management in river networks.

## Acknowledgments

A portion of this study was supported by JSPS KAKENHI Grant Numbers 26292181 and 19K20491, and Kuromatsunai Biodiversity Conservation Research Grant (2017). Dr. Izumi Washitani provided invaluable comments on earlier drafts of the manuscript. We thank the field assistance by Dr. Kosei Takahashi, Mr. Hitoshi Saito, Kengo Ebihara and Katsuya Iwabuchi. The all fish investigations were conducted with the permission of Hokkaido Prefecture. The authors have no competing interests.

